# phenopype: a phenotyping pipeline for Python

**DOI:** 10.1101/2021.03.17.435781

**Authors:** Moritz D. Lürig

## Abstract

1. Digital images are an intuitive way to capture, store and analyze organismal phenotypes. Many biologists are taking images to collect high-dimensional phenotypic information from specimens, which are key to investigating complex ecological, evolutionary and developmental phenomena, such as relationships between trait diversity and ecosystem function, multivariate natural selection, or developmental plasticity. As a consequence, images are being collected at ever increasing rates, but extraction of the contained phenotypic information still poses a veritable analytical bottleneck.
2. *phenopype* is a high throughput phenotyping pipeline for the programming language Python that aims at alleviating this bottleneck. The package facilitates immediate extraction of high dimensional phenotypic data from laboratory grade digital images with low levels of background noise and complexity. At the core, phenopype provides functionality for rapid signal processing based image preprocessing and segmentation, followed by trait extraction, visualization and numerous options for data export. The functionality is provided by wrapping low-level Python computer vision libraries (e.g. OpenCV) into accessible functions, facilitating their incorporation into scientific image analysis workflows. In addition, *phenopype* provides a project management ecosystem to further simplify rapid data collection and to increase reproducibility.
3. *phenopype* offers two different workflows that support users during different stages of scientific image analysis. The low-throughput workflow uses regular Python syntax and has greater flexibility at the cost of reproducibility, which is suitable for prototyping during the initial stage of a research project. The high-throughput workflow allows users to specify and store image-specific settings for analysis in human-readable YAML format, and then execute all functions in one step by means of an interactive parser. This approach facilitates rapid program-user interactions during batch processing, and greatly increases scientific reproducibility.
4. Overall, *phenopype* intends to make the features of powerful but technically involved low-level CV libraries available to biologists with little or no Python coding experience. Therefore, *phenopype* is aiming to augment, rather than replace the utility of existing Python CV libraries, allowing biologists to focus on rapid and reproducible data collection. Furthermore, image annotations produced by *phenopype* can be used as training data, thus presenting a stepping stone towards the application of deep learning architectures.

## 1 Introduction

Digital imaging has become one of the most widespread methods to study phenotypes in ecological and evolutionary research (Lürig et al., 2021; Pennekamp & Schtickzelle, 2013; Roeder et al., 2012). Produced by a myriad of devices such as digital cameras, scanners and microscopes, high resolution images capture a variety of internal, external or behavioral characteristics of organisms. In some cases, analysis of digital images replaced manual tasks, e.g. when using landmarks instead of manual calipers. Additionally, biologists are using images to draw a more complete picture of organismal phenotypes, e.g. through the analysis of colour (Ezray et al., 2019) and shape (Church et al., 2019), by tracing complex behavior (Dell et al., 2014; Manoukis & Collier, 2019), or by capturing microbial diversity (French et al., 2018; Zackrisson et al., 2016). Such data driven approaches bring the field closer to characterizing the phenome, i.e. the phenotype as a whole (Soulé, 1967), and help tracing causal links between phenotypes, genotypes and environmental conditions (Houle et al., 2010). In consequence, digital images are being collected at ever increasing rates, which has outpaced our ability to extract and analyze the phenotypic information contained within.

Computer vision (CV), the automated extraction and processing of information from digital images, has great potential to reduce image analysis as an analytical bottleneck in ecological and evolutionary research (Høye et al., 2021; Lürig et al., 2021; Weinstein, 2018). CV has already transformed data collection in various fields of research, such as medicine (Gao et al., 2018), remote sensing (Marmanis & Wegner, 2016), or material sciences (DeCost & Holm, 2015). In ecological and evolutionary research, however, software for CV is still only infrequently used to analyze image datasets (Lürig et al., 2021; Weinstein, 2018). Potential hurdles may lie in tradeoffs between flexibility and scalability on the one hand, and ease of use on the other hand. For instance, standalone toolboxes with a graphical user interface (GUI) such *ImageJ* (Schneider et al., 2012) or *ilastik* (Berg et al., 2019) are often preferred, because they offer a relatively broad range of CV-functions and machine learning capabilities while still being user friendly. However, batch analysis of larger image datasets is typically more convenient and efficient from within a high level programming language, such as Python, which has native file handling and array computing capabilities, as well as state-of-the art CV libraries. Yet, implementing an image analysis workflow from scratch often requires considerable technical knowledge, is specific to a given project or study system, and typically comes without even basic GUI functionality.

Here I introduce the Python package *phenopype*, which is a high throughput image analysis pipeline that allows ecologists and evolutionary biologists to extract high dimensional phenotypic data from digital images with little background noise. *Phenopype* resolves the aforementioned tradeoff by combining a rich (and further expandable) collection of CV functions implemented in Python for high throughput image analysis with basic GUI functionality. As a semi-manual method (Lürig et al., 2021) it allows users to step into an otherwise fully automatic workflow to correct for little differences between images and specimens. *phenopype* also comes with an efficient file and project management ecosystem to facilitate rapid data collection: all settings for image analysis are stored along with processed images and results, so that the acquired phenotypic information becomes highly reproducible. Overall, *phenopype* is aiming to augment, rather than replace the utility of existing Python CV libraries. Put differently, *phenopype* does neither intend to be an exhaustive library of granular image processing functions, like *OpenCV* or *scikit-image*, nor a purely GUI-based toolkit like *ImageJ*, but instead, it is a set of wrappers and convenient management tools that allows biologists to focus on rapid and reproducible data collection.

## 2 Implementation

### 2.1 Capabilities and limitations

*phenopype* is intended for the rapid extraction of trait data from digital images with low to moderate amounts of background noise, typically produced under laboratory conditions. Specifically, the specimens to be phenotyped should be photogrbaphed against a background with relatively constant colour and brightness (e.g., tray, petri-dish, microscope slide, scanner, etc.). This is because *phenopype*, in its current implementation, uses signal processing for segmentation; the process of splitting all pixels into foreground (objects of interest) and background (all remaining pixels). The advantages of this segmentation approach are that no GPU or training data are needed, and that all user interactions occur near-instantaneously (depending on image size and CPU power). Furthermore, users can filter out noise through various preprocessing steps (e.g. blur-kernels, morphology operations or masking), or correct flawed segmentation results with an interactive “paint-brush”. This makes *phenopype* especially suitable for tasks that involve the counting of specimens, shape measurements (up to 41 shape features), the extraction of colour intensities and texture from different colour channels (up to 120 texture features per channel), and movement recognition. For an overview of *phenopype*’s capabilities, refer to the examples shown on the project homepage (https://mluerig.github.io/phenopype/) and to section 4.

### 2.2 Package dependencies

*phenopype* (2.1.0) is implemented in Python 3.7. It relies on the Python Standard Library and several external dependencies, which are also downloaded during package installation (Fig. 1). Since images are represented as multidimensional arrays in Python, the *numpy* library (Harris et al., 2020) is needed to for basic array handling and computing capabilities. The OpenCV library ((Bradski, 2000), implemented through the *opencv-python* package) serves as the main backend for most image processing and analysis functionality (contained in the core modules), and for basic image I/O utility functions (Fig. 1A). Furthermore, *phenopype* uses the *pyradiomics* library (van Griethuysen et al., 2017) for the extraction of texture features. *phenopype* serves as a wrapper for *opencv-python* and *pyradiomics*, whose CV functions operate on a low level and thus require some basic checks and preparation steps, which are covered by the *phenopype* functions (Fig. 1B). The *ruamel*.*yaml* package is used to load and parse user modifications, and *watchdog*, a package that detects changes to files, is used to provide an interactive user experience while modifying YAML files.

**Figure 1.**
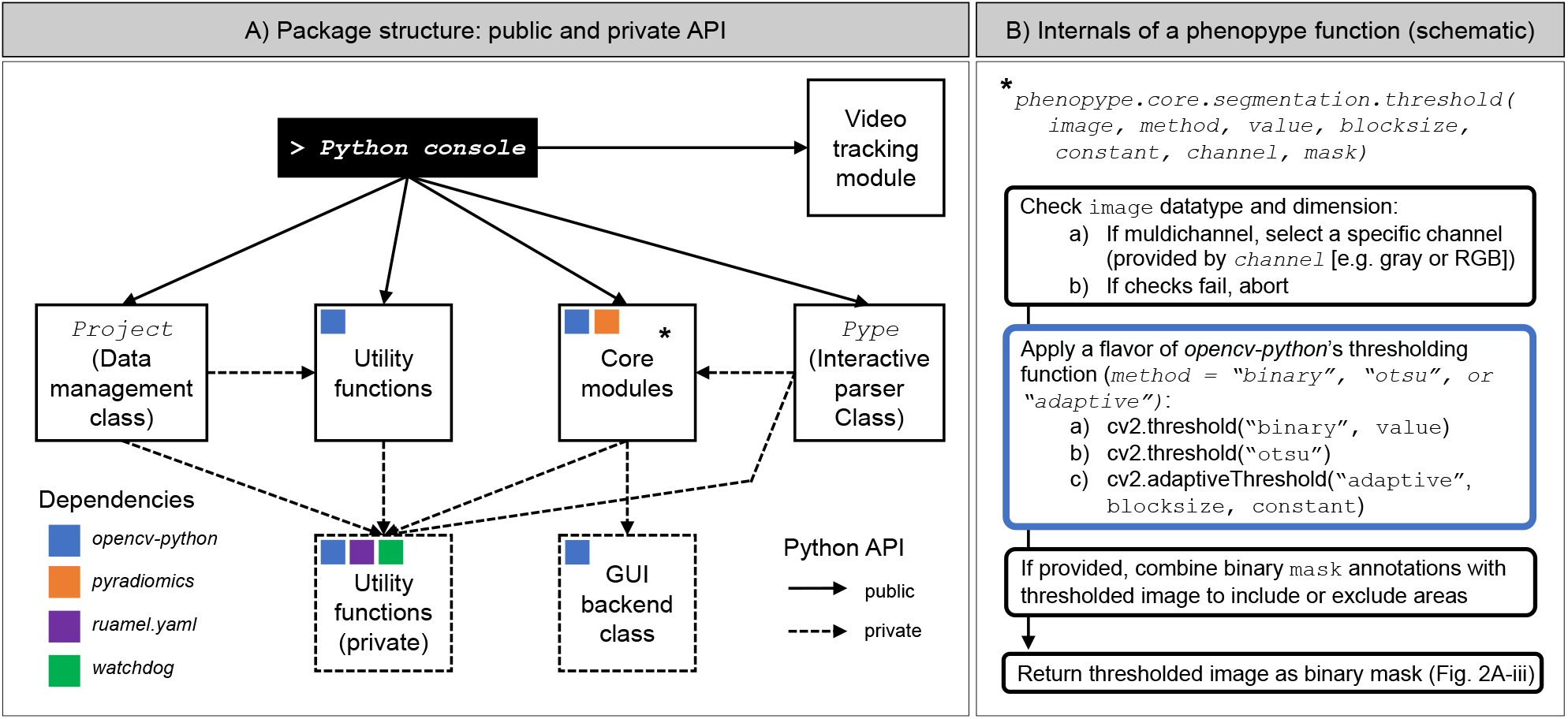
Schematic of the package design. A) The public API provides access to project management and utility functions (e.g. for loading and saving images), the core modules, which contain all image processing functions, and the *Pype* class, which is an interactive parser used in the high-throughput workflow (see section 3.2.2). B) *phenopype* does not supply its own technical basis for image analysis, but instead “wraps’’ existing low level computer vision functions, for example, from the *opencv-python* and *pyradiomics* libraries, into convenient functions that perform all necessary checks and conversions, thereby facilitating ease of use, as less coding is required from the user side. Here, a schematic for such a wrapper function (*threshold* function from the *segmentation* module) is given to exemplify how external libraries are integrated into *phenopype*.

### 2.3 Package API

The complete API (Application Programming Interface) is available online through the *phenopype* docs (https://mluerig.github.io/phenopype/), but a brief overview is given here (also see Fig. 2-3):

**Figure 2.**
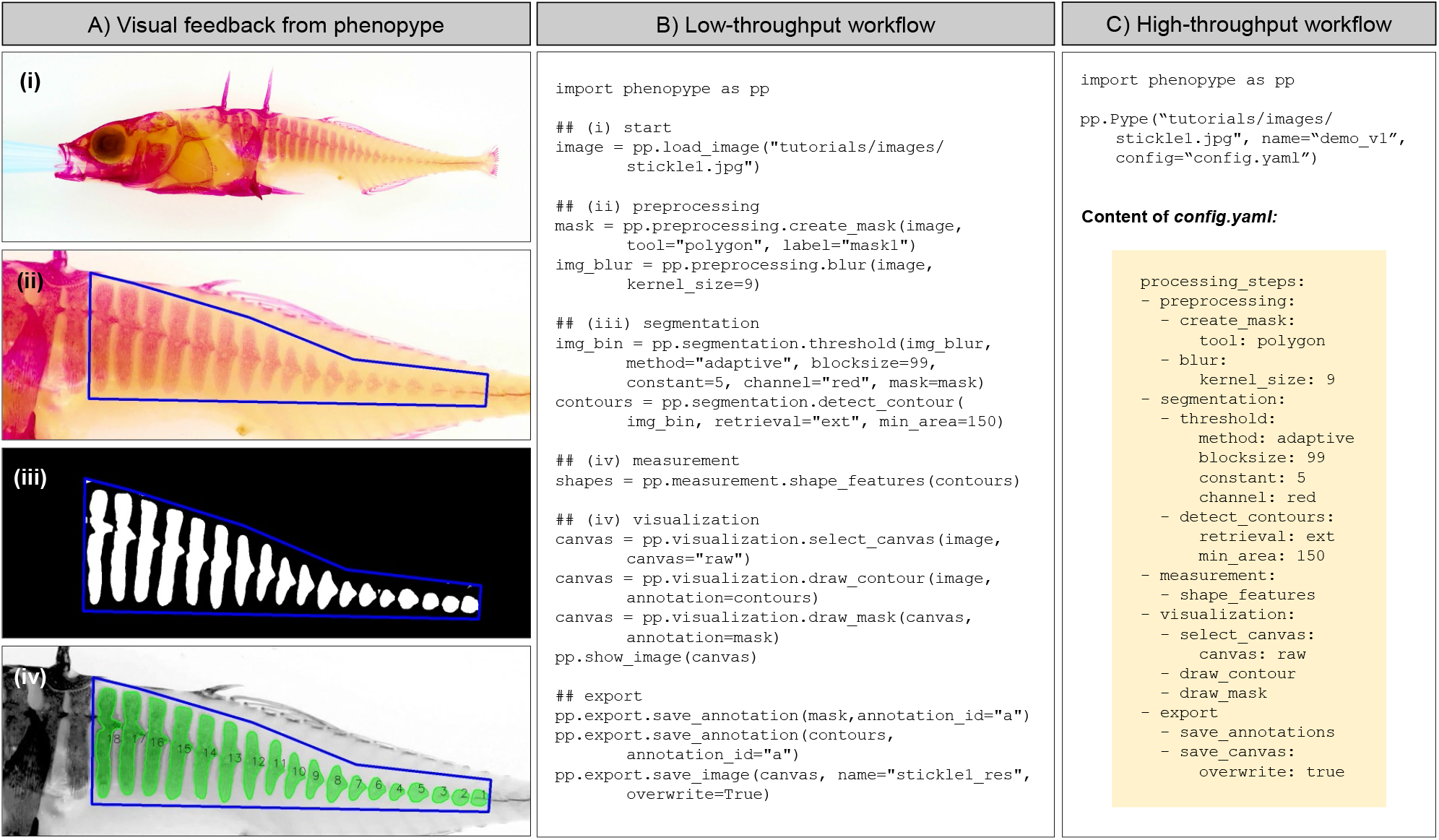
An exemplified computer vision task in *phenopype* to measure the number, area and shape of bony armor plating in a stained specimen of threespine stickleback (*Gasterosteus aculeatus*). Shown are visual feedback (A) from low-throughput (B) and high-throughput workflow (C), which use the same computer functions, but differ in the way these functions are called by the user: while in the low-throughput workflow all functions have to be explicitly coded in Python, the high-throughput routine parses the functions from human readable YAML files to facilitate rapid user interaction and increase reproducibility (see section 3.2 for details on both workflows). Image taken by Angelina Arquint.

**Figure 3.**
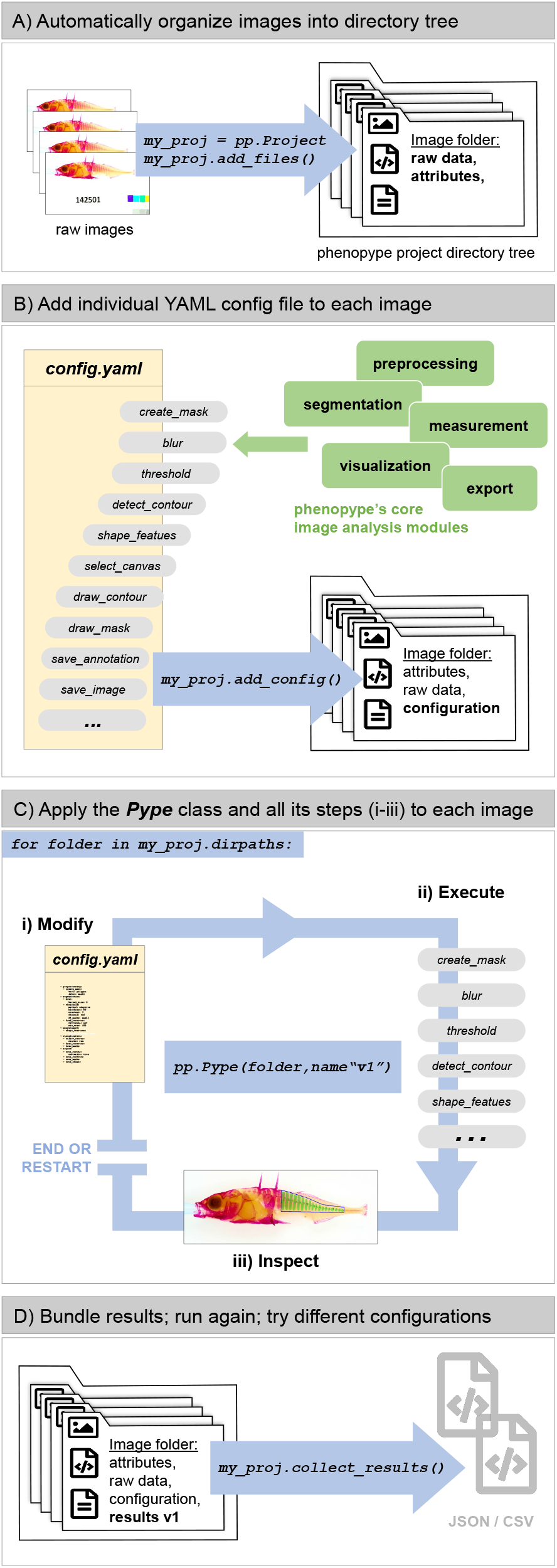
Flow chart of the high-throughput workflow in *phenopype*. A) First a *Project* class is initialized, yielding the class object *“my_proj”*. Then, using the class method *add_files*, a directory tree is be created where all relevant files are stored B) Next a configuration file in YAML format is created and stored within each folder of the *phenopype* directory tree using class method *add_config*. Users can choose between one of several configuration templates for different cases (e.g., counting landmarking, colour scoring, or shape measurement, and more). C) The *Pype* class in a *for* loop will trigger a series of events for each image directory provided by the loop generator: i) open the contained yaml configuration with the default OS text editor, ii) parse and execute the contained functions from top to bottom, iii) open a GUI window and show the processed image. Once the Pype class has finished executing all functions from the configuration file, users can decide to either modify the opened configuration file (e.g. either change function parameters or add new functions), which will trigger to run the Pype class again, or to close the GUI window, which will terminate the Pype class instance and save all results to the folder. By including the Pype class in a simple Python *for*-loop with all project folders, users can continue with this procedure throughout the entire project dataset, eliminating the need to manually open, close or save any of the images. D) When all images have been analyzed, the obtained results are obtained from each project folder with the class method *collect_results*, and store them in a new folder in the project root directory for further analysis in their favorite statistical environment.

- The project management class (*Project)* and its methods (e.g., *add_files, add_config, collect_results, etc*.) are intended to systematically select images on the harddrive and load them into an organized folder structure.
- Various utility functions for loading, inspecting and saving images (e.g., *load_image, show_image, save_image*, etc.).
- The five image analysis core modules (*preprocessing, segmentation, measurement, visualization, export*), with low-level CV tools from the *opencv-python* library wrapped into convenient functions (see Fig. 1B for a schematic). Some of these functions open a graphical user interface (GUI).
- The *Pype* class is an interactive parser, translating user instructions stored in YAML into Python code (see section 3.3 and Fig. 3C for details).

## 3 Getting started with phenopype

### 3.1 Installation and setup

To establish a Python 3.7 environment and to ensure the proper installation of all dependencies, the use of a Python environment manager is highly recommended (e.g., *Anaconda*). After setting up an environment, *phenopype* can be installed from the Python Package Index (PyPI; https://pypi.org/project/phenopype/) with *pip install phenopype*. Lastly, a text editor capable of opening YAML files (without locking them) needs to be installed and selected as the default program.

### 3.2 Working with phenopype

In *phenopype*, users can choose between “low-throughput” and “high-throughput” workflows, each of which accommodate the different stages in the process of scientific image analysis. Early during an ongoing experiment or a survey, when images are first evaluated on the computer screen, scientific image analysis can be a highly iterative process that may require frequent user input to preprocess and segment images, and then evaluate the obtained results. This may be faster and more intuitive with the low-throughput workflow, which allows inspection of the modified arrays and annotations in Python. Later, when the best functions and appropriate settings are found and rapid data collection has priority, image analysis should be seamless and with minimal user input for maximal throughput, but also with maximal scientific reproducibility, which is ensured with the high-throughput workflow.

#### 3.2.1 Low-throughput workflow

In the low-throughput workflow, images are loaded as arrays using the utility functions, and then processed and analyzed using CV functions from the five core modules are called in Python (Fig. 2B). This allows users to interact with arrays directly, which is providing a good sense of which function is doing what and likely a good starting point for people with no prior exposure to the concepts of computer vision. However, this also means that settings need to be modified for each image separately, which makes this workflow inconvenient for larger datasets.

#### 3.2.2 High-throughput workflow

Here, instead of coding each function for each image, the entire workflow is stored in human readable YAML format and can be called by a special interactive parser class (*Pype* class; Fig. 2C, Fig. 3). This dramatically reduces user input, and is thus suitable for the processing of larger datasets (100s - 1000s of images) using a simple *for* loop (Fig. 3C). To further support batch processing, *phenopype* provides convenient project management functions to store and organize raw image data, YAML configurations, and results in a directory tree (*Project* class; Fig. 3).

#### 3.2.3 Video analysis

*phenopype* also includes a video analysis module which allows segmentation of video frames using foreground-background subtraction algorithms. Such algorithms first determine what constitutes the background by detecting static pixels over a range of frames, and then determine foreground by identifying pixels of objects that deviate from the background model through movement. The video module, although fully functional and with complete documentation, is currently only weakly integrated into the different *phenopype* workflows, and is listed here only for completeness.

## 4 Feature demonstration

*phenopype* can be used to extract a wide array of organismal traits, particularly those related to external shape and texture, and it has already proven its effectiveness by producing large amounts of phenotypic data for two peer reviewed publications (Lürig et al., 2019; Lürig & Matthews, 2021). Here I demonstrate some of *phenopype*’s key features and provide some examples of different types of trait data that can be extracted. The following paragraphs correspond to the panels in Fig. 4. More extensive examples with code can be found in the online documentation (https://mluerig.github.io/phenopype/):

**Figure 4.**
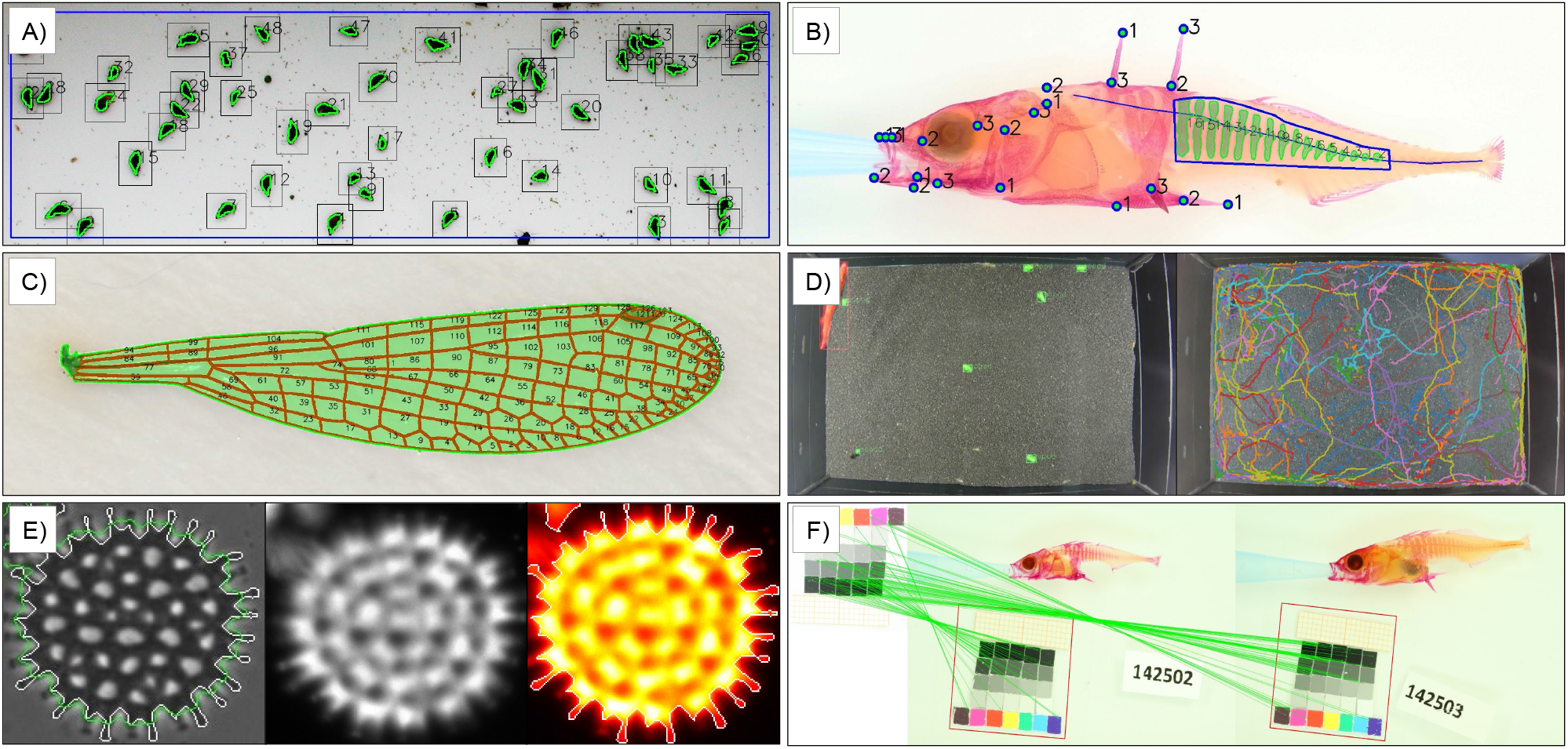
Demonstration of major *phenopype* features (the panels correspond to the paragraphs in section 4). A) Counting and measuring freshwater snails (*Potamopyrgus antipodarum;* image by Jukka Jokela). B) Landmarking and armor plate measurement in threespine stickleback (*Gasterosteus aculeatus*; image by Dominique Stalder). C) Quantifying venation patterns in damselfly wings (*Ischnura elegans*; image by Masahito Tsuboi). D) Tracking predator prey interactions between stickleback (*G. aculeatus)* and freshwater isopods (*Asellus aquaticus;* video shot by Kim Kaltenbach). E) Extracting shape and fluorescence traits from phytoplankton cells (*Pediastrum sp*.; image by Irene Gallego). F) Automatic detection of reference cards in images using *phenopype* (image by Dominique Stalder).

A. *Counting snails:* Counting small organisms in the laboratory is a task that has good potential to be solved automatically with CV. Here, *phenopype* was used to count freshwater snails (*Potamopyrgus antipodarum*), and measure their body size, shape and pigmentation. This was achieved through consecutive use of threshold (to segment black snails from white background), and watershed algorithms (to separate snails that are touching each other). Small fecal pellets are filtered from the detected foreground (the green contours) using blur kernels and a size cutoff. Corrections for the gradient of brightness in the image are conducted by comparing each specimen to the background within its bounding box.
B. *Stickleback landmarks and armor plating*: Functional morphology of organisms is often measured by placing landmarks at specific points, but variation in continuous phenotypic traits like shape or area are difficult to quantify manually because they are too complex or have no underlying assumption of homology. In this example, the traits of interest are contained in 22 landmarks that were set with *phenopype* across the anterior half of a threespine stickleback (*Gasterosteus aculeatus*), which was stained with alizarin red. In addition, the number, area and extension of armor plating was measured as a continuous trait: first, a mask was set around the posterior region that contains the plates, then the red colour channel of the image, which contained the highest signal-to-noise ratio, was thresholded.
C. *Venation patterns in damselfly wings*: Here, *phenopype* was used to measure the spacing, angles, and connectivities within the wing tissue of an adult damselfly (*Ischnura elegans*). Because the venation within the wing has an extremely high signal-to-noise ratio, a single thresholding algorithm could be used without further preprocessing. The full hierarchy of foreground contours was resolved: the outer perimeter of the whole wing (in green) and all domains within the wing (in red).
D. *Isopod movement in response to predation:* In this example, the goal was to track the movement of freshwater isopods (*Asellus aquaticus*) in response to presence, absence and activity of a predator (*G. aculeatus*). The output of the *phenopype*’s video tracking module is a time series of coordinates that correspond to the trajectories of predator and prey. In addition, the foreground-background subtraction algorithm produces a binary segmentation mask for every frame. This allows the detection of pigmentation and body size of every isopod, and thus the sequence in which the fish consumed differently pigmented and sized specimens.
E. *Phytoplankton cell shape and fluorescence:* A plate reader produced bright-field and fluorescence emission images of freshwater phytoplankton communities (shown here is *Pediastrum sp*.). Here the goal was to measure the shape (cell morphology) and texture (fluorescence intensity) of cells. This was done in a two step process: first, the brightfield image was thresholded to detect the boundary contours of the cell. Then, the contours were used as a stencil on images of different fluorescence channels that capture variation in three pigments (chlorophyll, phycoerythrin, phycocyanin), measuring intensities within the contour of the cell.
F. *Automatic reference card detection*: Even in highly standardized laboratory environments, there are often slight variations in camera exposure and zoom that may affect the outcome of measurements and extracted traits. To automate the time consuming task of correcting for size ratio and brightness variation, *phenopype* provides a suite of performant image registration algorithms (Tareen & Saleem, 2018) to detect reference cards within images, and automatically correct size and colour histograms of that image. This can be done on a project wide scale, where first a template is created, and then detected in every following image.

## 5 Discussion and outlook

With *phenopype* I have presented a novel approach for rapid, reproducible and user-friendly extraction of phenotypic data from digital images. Its main innovation lies in making low level CV-functions more accessible through wrappings and markup language that is both human and machine readable: by using YAML configuration files instead of Python code users can assemble functions in manifold ways to address the specifics of their study organism, while maintaining cross-platform compatibility and achieving complete reproducibility. Together with a simple GUI and a convenient project management ecosystem, this approach provides a seamless workflow for larger image datasets. Initially, users may take some time to learn and understand the approach. Here the low-throughput workflow can serve as an easy introduction to computer vision in Python. In the long run, the initial time investment will pay off, as tedious and repetitive tasks are eliminated with the high-throughput workflow, which should save a lot of time during image analysis and project organization.

*phenopype* provides CV functions that are based on signal processing, which work instantaneously (i.e. they don’t require any training), have lower technical hurdles (no GPU required), and are less complex to implement, but are prone to higher levels of background noise and convolution. Therefore, the package is intended as a pipeline for scientific photography dedicated to phenotyping, where by default great care is taken to minimize levels of background noise. While for this purpose machine learning is typically not needed to achieve good segmentation, it moreover is often not even possible, because training data, sufficient time, or GPU-hardware appropriate for deep learning are lacking (Lürig et al., 2021). For cases without these constraints, with very large datasets, or with high levels of noise, recent innovative Python machine learning toolkits like *ml-morph* (Porto & Voje, 2020) or *sashimi* (Schwartz & Alfaro, 2021) are recommended alternatives.

Regardless of potential constraints or scope of the research, *phenopype*, may serve as a stepping stone towards such machine learning approaches, because it produces foreground masks as a “byproduct” and thus provides labelled training datasets, which are an increasing bottleneck for deep learning approaches (Sun et al., 2017). In the long term and with the help and feedback of the CV community, a deep learning workflow could even be integrated to *phenopype*: on the one hand, Python, specifically the *OpenCV* library that *phenopype* is using as a backend, already comes with a suite of state-of-the-art deep learning implementations, which are frequently updated with the most cutting edge algorithms (such as Mask-RCNN (Abdulla, 2017)). On the other hand, *phenopype’s* modular architecture facilitates the inclusion of current and future advances within the highly dynamic field of scientific CV.

## Data availability statement

A snapshot of *phenopype* in version 2.1.0, as well as the full documentation, including installation instructions, API reference and tutorials can be found on Zenodo (Lürig, 2021) and on GitHub (master branch: https://github.com/mluerig/phenopype; documentation branch: https://mluerig.github.io/phenopype/).

## Acknowledgements

I conceived *phenopype* in its current form during a lab retreat in Vna (Graubünden, Switzerland) that was organized and funded by Jukka Jokela. Its implementation was made possible by Blake Matthews and the Eawag directorate (Discretionary Funding Grant No. 5221.00492.013.11). Additional funding came from the Swiss National Science Foundation through an Early Postdoc.Mobility Fellowship (Grant No. P2EZP3_191804) and from the European Union’s Horizon 2020 research and innovation programme through a Marie Skłodowska-Curie IF (Grant No. 898932). I thank Kim Kaltenbach for being a patient alpha tester, and Cam Hudson, Ryan Greenway, Andres Grolimund, Nare Ngoepe, Anja Merz and Irene Gallego for being helpful beta testers. I would also express my sincere gratitude to Arthur Porto and Seth Donoughe whose comprehensive review for the *pyOpenSci* consortium greatly improved the presentation and documentation of the package. Two anonymous reviewers provided very constructive feedback during review of the manuscript. The stickleback image for *phenopype’s* logo was taken by Angelina Arquint. Finally, this package may not have come into existence without Matt McGee, who encouraged me to learn Python and use it for computer vision.

## References

Abdulla, W. (2017). Mask r-cnn for object detection and instance segmentation on keras and tensorflow. https://github.com/matterport/Mask_RCNN

Berg, S., Kutra, D., Kroeger, T., Straehle, C. N., Kausler, B. X., Haubold, C., Schiegg, M., Ales, J., Beier, T., Rudy, M., Eren, K., Cervantes, J. I., Xu, B., Beuttenmueller, F., Wolny, A., Zhang, C., Koethe, U., Hamprecht, F. A., & Kreshuk, A. (2019). ilastik: interactive machine learning for (bio)image analysis. Nature Methods, 16(12), 1226–1232. https://doi.org/10.1038/s41592-019-0582-9

Bradski, G. (2000). The opencv library. Doctor Dobbs Journal, 25(11), 120–126.

Church, S. H., Donoughe, S., de Medeiros, B. A. S., & Extavour, C. G. (2019). Insect egg size and shape evolve with ecology but not developmental rate [Review of Insect egg size and shape evolve with ecology but not developmental rate]. Nature, 571(7763), 58–62. https://doi.org/10.1038/s41586-019-1302-4

DeCost, B. L., & Holm, E. A. (2015). A computer vision approach for automated analysis and classification of microstructural image data. Computational Materials Science, 110, 126–133. https://doi.org/10.1016/j.commatsci.2015.08.011

Dell, A. I., Bender, J. A., Branson, K., Couzin, I. D., de Polavieja, G. G., Noldus, L. P. J. J., Pérez-Escudero, A., Perona, P., Straw, A. D., Wikelski, M., & Brose, U. (2014). Automated image-based tracking and its application in ecology. Trends in Ecology & Evolution, 29(7), 417–428. https://doi.org/10.1016/j.tree.2014.05.004

Ezray, B. D., Wham, D. C., Hill, C. E., & Hines, H. M. (2019). Unsupervised machine learning reveals mimicry complexes in bumblebees occur along a perceptual continuum. Proceedings. Biological Sciences / The Royal Society, 286(1910), 20191501. https://doi.org/10.1098/rspb.2019.1501

French, S., Coutts, B. E., & Brown, E. D. (2018). Open-Source High-Throughput Phenomics of Bacterial Promoter-Reporter Strains. Cell Systems, 7(3), 339–346.e3. https://doi.org/10.1016/j.cels.2018.07.004

Gao, J., Yang, Y., Lin, P., & Park, D. S. (2018). Computer Vision in Healthcare Applications. Journal of Healthcare Engineering, 2018, 5157020. https://doi.org/10.1155/2018/5157020

Harris, C. R., Millman, K. J., van der Walt, S. J., Gommers, R., Virtanen, P., Cournapeau, D., Wieser, E., Taylor, J., Berg, S., Smith, N. J., Kern, R., Picus, M., Hoyer, S., van Kerkwijk, M. H., Brett, M., Haldane, A., Del Río, J. F., Wiebe, M., Peterson, P., … Oliphant, T. E. (2020). Array programming with NumPy. Nature, 585(7825), 357–362. https://doi.org/10.1038/s41586-020-2649-2

Houle, D., Govindaraju, D. R., & Omholt, S. (2010). Phenomics: the next challenge. Nature Reviews. Genetics, 11(12), 855–866. https://doi.org/10.1038/nrg2897

Høye, T. T., Ärje, J., Bjerge, K., Hansen, O. L. P., Iosifidis, A., Leese, F., Mann, H. M. R., Meissner, K., Melvad, C., & Raitoharju, J. (2021). Deep learning and computer vision will transform entomology. Proceedings of the National Academy of Sciences of the United States of America, 118(2). https://doi.org/10.1073/pnas.2002545117

Lürig, M. D. (2021). Data from: phenopype: a phenotyping pipeline for Python. https://doi.org/10.5281/zenodo.4609990

Lürig, M. D., Best, R. J., Svitok, M., Jokela, J., & Matthews, B. (2019). The role of plasticity in the evolution of cryptic pigmentation in a freshwater isopod. The Journal of Animal Ecology, 88(4), 612–623. https://doi.org/10.1111/1365-2656.12950

Lürig, M. D., Donoughe, S., Svensson, E. I., Porto, A., & Tsuboi, M. (2021). Computer Vision, Machine Learning, and the Promise of Phenomics in Ecology and Evolutionary Biology. Frontiers in Ecology and Evolution, 9, 148. https://doi.org/10.3389/fevo.2021.642774

Lürig, M. D., & Matthews, B. (2021). Dietary-based developmental plasticity affects juvenile survival in an aquatic detritivore. Proceedings of the Royal Society B: Biological Sciences, 288(1945), 20203136. https://doi.org/10.1098/rspb.2020.3136

Manoukis, N. C., & Collier, T. C. (2019). Computer Vision to Enhance Behavioral Research on Insects. Annals of the Entomological Society of America. https://doi.org/10.1093/aesa/say062

Marmanis, D., & Wegner, J. D. (2016). Semantic segmentation of aerial images with an ensemble of CNNs. Annals of the …. https://www.isprs-ann-photogramm-remote-sens-spatial-inf-sci.net/III-3/473/2016/isprs-annals-III-3-473-2016.pdf

Pennekamp, F., & Schtickzelle, N. (2013). Implementing image analysis in laboratory-based experimental systems for ecology and evolution: a hands-on guide. Methods in Ecology and Evolution / British Ecological Society, 4(5), 483–492. https://doi.org/10.1111/2041-210X.12036

Porto, A., & Voje, K. L. (2020). ML‐morph: A fast, accurate and general approach for automated detection and landmarking of biological structures in images. Methods in Ecology and Evolution / British Ecological Society, 2041-210X.13373. https://doi.org/10.1111/2041-210x.13373

Roeder, A. H. K., Cunha, A., Burl, M. C., & Meyerowitz, E. M. (2012). A computational image analysis glossary for biologists. Development, 139(17), 3071–3080. https://doi.org/10.1242/dev.076414

Schneider, C. A., Rasband, W. S., & Eliceiri, K. W. (2012). NIH Image to ImageJ: 25 years of image analysis. Nature Methods, 9(7), 671–675. https://doi.org/10.1038/nmeth.2089

Schwartz, S. T., & Alfaro, M. E. (2021). Sashimi : A toolkit for facilitating high‐throughput organismal image segmentation using deep learning. Methods in Ecology and Evolution / British Ecological Society, 2041-210X.13712. https://doi.org/10.1111/2041-210x.13712

Soulé, M. (1967). PHENETICS OF NATURAL POPULATIONS I. PHENETIC RELATIONSHIPS OF INSULAR POPULATIONS OF THE SIDE-BLOTCHED LIZARD. Evolution; International Journal of Organic Evolution, 21(3), 584–591. https://doi.org/10.1111/j.1558-5646.1967.tb03413.x

Sun, C., Shrivastava, A., Singh, S., & Gupta, A. (2017). Revisiting Unreasonable Effectiveness of Data in Deep Learning Era. In arXiv [cs.CV]. arXiv. http://arxiv.org/abs/1707.02968

Tareen, S. A. K., & Saleem, Z. (2018, March 4). A comparative analysis of SIFT, SURF, KAZE, AKAZE, ORB, and BRISK. ResearchGate. 2018 International Conference on Computing, Mathematics and Engineering Technologies – iCoMET 2018. https://doi.org/10.1109/ICOMET.2018.8346440

van Griethuysen, J. J. M., Fedorov, A., Parmar, C., Hosny, A., Aucoin, N., Narayan, V., Beets-Tan, R. G. H., Fillion-Robin, J.-C., Pieper, S., & Aerts, H. J. W. L. (2017). Computational Radiomics System to Decode the Radiographic Phenotype. Cancer Research, 77(21), e104–e107. https://doi.org/10.1158/0008-5472.CAN-17-0339

Weinstein, B. G. (2018). A computer vision for animal ecology. The Journal of Animal Ecology, 87(3), 533–545. https://doi.org/10.1111/1365-2656.12780

Zackrisson, M., Hallin, J., Ottosson, L.-G., Dahl, P., Fernandez-Parada, E., Ländström, E., Fernandez-Ricaud, L., Kaferle, P., Skyman, A., Stenberg, S., Omholt, S., Petrovič, U., Warringer, J., & Blomberg, A. (2016). Scan-o-matic: High-Resolution Microbial Phenomics at a Massive Scale. G3, 6(9), 3003–3014. https://doi.org/10.1534/g3.116.032342

